# *AtERF60,* negatively regulates *ABR1* to modulate basal resistance response in Arabidopsis

**DOI:** 10.1101/2021.12.29.474403

**Authors:** Ashutosh Joshi, Ujjal J Phukan, Neeti Singh, Gajendra Singh Jeena, Sunil Kumar, Durgesh Kumar Pandey, Alok Pandey, Vineeta Tripathi, Rakesh Kumar Shukla

## Abstract

APETALA2 (AP2)/ERF family transcription factors (TFs) contribute an important function against various external cues. Our study reports *AtERF60*, an AP2/ERF TF, which plays a key role in regulating the *ABR1* gene, leading to an altered basal resistance response in Arabidopsis. *AtERF60* is induced in response to drought, salt, abscisic acid (ABA), salicylic acid (SA), and bacterial pathogen *Pst*DC3000 infection. *AtERF60* interacts with DEHYDRATION RESPONSE ELEMENTS and GCC box, indicating its ability to regulate multiple responses. The overexpressing lines of *AtERF60* demonstrated increased resistance to *Pst*DC3000 infection, whereas *erf60* mutant lines showed increased susceptibility. Complementation of the *erf60* mutant background exhibits no significant difference towards *Pst*DC3000 infection compared with the WT Col-0. Microarray and qRT-PCR analysis of overexpression and mutant lines indicated that *AtERF60* regulates stress-inducible genes. The induction of these differentially expressed transcripts was significantly increased in *erf60* mutant lines, whereas it was reversed in wild-type lines when *AtERF60* was complemented. *ABR1* was one of the differentially expressed transcripts, and we discovered that *AtERF60* interacts with the DRE *cis*-elements in the *ABR1* promoter. Further, qRT-PCR expression analysis after infection with *Pst*DC3000 in *AtERF60-OX*, *erf60* and its complementation background suggest that the mutation in *AtERF60* upregulates *ABR1* activity, leading to the enhanced susceptibility towards *Pst*DC3000. Conversely, *AtERF60* overexpression suppresses *ABR1* activity, strengthening the basal resistance responses of Arabidopsis.

## Introduction

Plants face multiple environmental stresses, including biotic and abiotic stresses, which disturb their normal growth and development, leading to massive agricultural losses (He *et al*., 2018). To adapt to these environmental conditions, plants have developed various evolutionary mechanisms governed by a complex regulatory network. Transcription factors regulate different interconnected and diverse signalling cascades by interacting with *cis*-elements present in their promoters (Liu *et al*., 2014). AP2/ERFs are one of the most important transcription factor families that appeared as a key regulator of a large cluster of downstream target genes involved in various stress responses (Phukan *et al*., 2017). The AP2/ETHYLENE RESPONSIVE ELEMENT BINDING FACTOR (EREB) domain of AP2/ERFs is made up of conserved 40–70 amino acids that act as a DNA binding domain (Nakano *et al*., 2006). Based on the DNA binding domain, AP2/ERFs are classified into three groups: first is ERF (single ERF domain, contains subgroup I to X, VI-L and Xb-L), second is AP2 (tandem copies of two AP2 domains and few of them are with single AP2 domain), and third is RAV (ERF domain associated with a B3 DNA-binding domain). Soloist is a protein that contains an ERF domain; however, its sequence and gene structure strongly diverge from ERF TFs (Shigyo and Ito, 2004, Nakano *et al*., 2006, Swaminathan *et al*., 2008, Zhuang *et al*., 2008; Licausi *et al*., 2010, Licausi *et al*., 2013). A member of AP2/DREB type TF, RAP2.4 was induced in response to salt and drought stress (Feng *et al*., 2005). *WOUND INDUCED DEDIFFERENTIATION 1–4* (*WIND1-4*), a member of RAP2.2, are induced on wounding and play important roles in callus formation at wounded sites (Iwase *et al*., 2011). These proteins integrate multiple phytohormone signals and regulate abiotic and biotic stress responses (Gutterson and Reuber, 2004). ERF binds with the ETHYLENE-RESPONSE ELEMENT (ERE) or GCC-box to provide biotic stress resistance (Franco-Zorrilla *et al*., 2014).

Different properties of AP2/ERFs, including the DNA binding properties and induction upon various stresses, implement these TFs to coordinate multiple responses and participate in regulatory processes (Abiri *et al*., 2017). It is evident that AP2/ERFs and various phytohormones such as ethylene, Methyl jasmonate (MeJA), ABA, SA act in concert to regulate various plant processes (Licausi *et al*., 2013). The coordinated action of ABA and different abiotic stresses leads to the activation of several stress-inducible genes and DREB2s (Lee *et al*., 2016). AP2/ERFs such as *ABA-INSENSITIVE 4* (*ABI4*) and *CBFA* (*CCAAT binding factor A*) are ABA-responsive and help to activate the ABA-dependent/independent stress-responsive genes (Zhang *et al*., 2013). AP2/ERF mutants with altered hormone sensitivity and abiotic stress responses have been identified and studied, mentioning this family of TFs as an important candidate for studying the interactions between plant hormones and abiotic stresses.

The ERF subfamily protein is usually conscious of pathogen attack and contributes to plant immunity. ERF subfamily protein’s overexpression is generally related to altered disease resistance phenotypes in plants (Onate-Sanchez *et al*., 2007; Zhang *et al*., 2016; Zheng *et al*., 2019). Overexpression of rice *OsEREBP1* activates ABA and JA signalling pathways, eventually resulting in enhanced tolerance under biotic and abiotic conditions (Jisha *et al*., 2015). ABA regulates various important agronomical traits of plant development and physiology, such as seed dormancy and maturation, along with responses to several environmental stress factors, such as salinity, drought, and cold stress mediated through different components of ABA signalling (Himmelbach *et al*., 2003; Sah *et al*., 2016). During the investigation of gene expression regulated with ABA, an AP2-domain containing a protein known as *ABR1* that regulates ABA responses was identified (Pandey *et al*., 2005). Later, it was studied that *ABR1* (group X of AP2/ERF) acts as a transcriptional activator and is involved in the wounding response (Baumler *et al*., 2019). *ABR1* is an important molecule thought to play a functional role in the host-pathogen interface and acts as a susceptibility core that binds with different effector molecules of *Pseudomonas syringae* (Schreiber *et al*., 2021). Despite the importance of *ABR1* in regulating biotic stress responses, its transcriptional regulator is unknown. In our study, we have identified and characterized *AtERF60* (Closest Arabidopsis homologue of our previously characterized protein MaRAP2-4 from *Mentha arvensis*) (Phukan *et al*., 2018). We investigated that *AtERF60* regulates the *ABR1* gene by binding with their promoter and modulates basal stress response in Arabidopsis.

## Materials and Methods

### Plant growth conditions and stress treatment

Arabidopsis Col-0 ecotype was utilized in this study. The plants were grown in the Conviron growth chamber model A-1000 as described in Phukan *et al*. (2018). The T-DNA insertion mutant lines were obtained from the Arabidopsis Biological Resource Centre (ABRC) and screened for homozygous lines. For growing, the Arabidopsis seeds were surface sterilized (3% sodium hypochlorite solution), stratified at 4^0^C for 96 hours, and then transferred to the growth chamber under controlled conditions with photoperiod (16:8 h light-dark) at 22°C and 60% relative humidity. MeJA treatment was provided by spraying a 100 μM solution made in dimethyl sulfoxide (DMSO) and Triton-X. Control plants were sprayed with a solution containing only DMSO and Triton-X. For SA and ABA treatment, plants were sprayed with 100 μM of SA and ABA, while control plants were sprayed with water. The aerial part, including the stem and leaves, was punctured with a needle to avoid major injuries. To avoid contamination, treated samples were obtained at various periods and cleaned thoroughly with sterile water. Two independent lines of Salk_134892 were utilized in this study.

### Expression analysis and phylogenetic analysis

Total RNA was isolated from plant samples utilizing RNeasy Plant Mini Kit (Qiagen), and cDNA was prepared utilizing a high-capacity cDNA reverse transcription kit (Applied Biosystems). Total cDNA was checked for quality by PCR using the control primers provided in the kit. The qRT-PCR (Applied biosystems 7900-HT Fast Real-Time PCR) was performed to determine the expression level of transcripts utilizing the SYBR Green PCR master mix kit (Takara). The unique region of transcripts was selected for designing qRT-PCR primers and mentioned in Table S3. Actin and ubiquitin were used as endogenous controls for normalizing gene expression. The 2^-ΔΔCt^ method was utilized to calculate the relative expression of genes. The protein sequences of all the AP2/ERF family members related to Arabidopsis were downloaded from the TAIR database. The sequence alignment was performed by using clustal omega online program using default parameters (https://www.ebi.ac.uk/Tools/msa/clustalo/), and the phylogenetic tree was constructed using MEGAXI using the neighbour-joining method, with 1000 bootstrap replicates.

### Cloning experiments, generation of transgenic lines, and mutant screening

The *AtERF60* was amplified using gene-specific primers and cloned in the PTZ57R/T vector. To analyze the in vitro interaction of protein and DNA, *AtERF60* was cloned in pGEX-4-T2 expression vector fused with GST. The positive clones were confirmed by PCR and digestion with primer-specific restriction enzymes (*BamHI* and *Xho*I) (Fig. S2a). The resulting positive construct was transformed in Lon and OmpT protease deficient *E. coli BL21* (DE3) strain for expression in the bacterial system. To study the functional role of *AtERF60* in planta, we cloned it in the pBI121 binary vector downstream of the CaMV35S promoter. The positive clones were confirmed by PCR, sequencing and final digestion utilizing the specific restriction enzymes (*BamHI* and *Xba*I) (Fig. S3a). The *AtERF60* transgenic lines in Arabidopsis were raised using the *Agrobacterium*-mediated gene transformation (Fig. S3b). Four independent transgenic lines containing *AtERF60* were generated, and two lines showing similar responses were used in the study. Transgenic lines in Arabidopsis were confirmed by using genomic DNA PCR using pBI-121 nptII (Kan^R^) specific primers. The PCR amplification ensures the successful integration of the desired gene (Fig. S3c,d). To determine the functional role of the *AtERF60*, we used T-DNA mutant lines of *AtERF60* (SALK_138492). The *erf60* mutants were screened using genomic DNA PCR with specific primers designed from the left border of T-DNA (Fig. S4). Two independent complementation lines were generated by transforming *AtERF60* in *erf60* mutant background, followed by screening on kanamycin for three generations to obtain the homozygous lines.

### Electrophoretic mobility shift assay

To study the in vitro protein-DNA interaction, *AtERF60* was cloned in the bacterial expression vector pGEX-4-T2. The open reading frame was continued with GST without disturbing the amino acid sequences. The cloned construct was then transformed in Lon and OmpT protease deficient *E. coli BL21* (DE3) strain and induced with 0.4 mM IPTG at 37°C for 5 h. The recombinant fusion protein was purified with Glutathione Sepharose beads (Sigma). Definite probes were designed for different *cis*-elements and their mutated versions (Table S3). EMSA followed the manufacturer’s protocol mentioned in the 2^nd^ generation DIG gel shift kit (Roche).

### Microarray analysis

The complete RNA was extracted from the plant samples (WT, *ERF60*-OX, and *erf60* mutant) under controlled conditions utilizing the RNeasy Plant Mini Kit (Qiagen) following the manufacturer’s instructions. The Arabidopsis GXP 4×44K AMADID slide was used to hybridize the RNA, which was further aided and labelled with Cy3-CTP. The microarray was performed and scanned at 535 nm, and the images were analyzed using Agilent Feature Extraction software (v10.7) to calculate signal and background strength. The images obtained from the microarray were cleaned and are uniformly intense, with very little background noise. Gene-Spring GX 12.6 software was utilized for statistical significance and normalization. The fold induction values obtained from two different lines were normalized to a single fold induction. The student’s t-test was used to correct P-values for the downregulation and upregulation of genes in experimental and biological replicates. The P-value cut-off for gene up- and downregulation was set to <0.05.

### Bacterial pathogen assays with *Pseudomonas syringae* pv. tomato DC3000 (*Pst*DC3000)

Bacterial pathogen *Pst*DC3000 was grown at 28°C overnight on LB media containing 50μg/ml rifampicin. For infiltration, bacteria were re-suspended in 10 mM MgCl_2_ to obtain OD_600_ = 0.2. The leaves of 5-week-old WT, *AtERF60-*OX, *erf60* mutant, and *AtERF60*-complementation Arabidopsis plants were syringe infiltrated with *Pst*DC3000. Disease symptoms were observed and photographed at regular intervals. Bacterial populations [Log (CFU/cm^2^)] in leaf tissues of WT, *AtERF60-*OX, *erf60* and *AtERF60*-complementation plants at 0, 2, and 4 dpi, were determined according to Katagiri *et al*. (2002). Four leaf discs of equal area (0.5 cm2) were taken from each WT, *AtERF60*-OX, *erf60* and *AtERF60*-complementation plants inoculated with *Pst*DC3000 at 4 dpi for electrolyte leakage assay. These leaf discs were then agitated in a tube containing 10 ml milli-Q water for 3 h. The conductivity was measured using the electrical conductivity meter for each sample (HORIBA Scientific, F74BW). Afterwards, leaf discs for each sample were autoclaved to release the total ions, and the conductivity corresponding to total ions was measured. Electrolyte leakage values (conductivity value at 3 h) were presented as the percentage relative to total ions.

Pathogen colonization in Arabidopsis plants was visualized using a Zeiss confocal laser-scanning microscope (LSM-880). We used GFP-tagged *Pst*DC3000 for this assay. Confocal images of leaf samples of WT, *AtERF60-*OX, *erf60* and *AtERF60*-complementation plants inoculated with GFP-tagged *PstD*C3000 were taken at 2 dpi. GFP acquisition was performed at 488 nm excitation with emission collection at 493-598 nm. Image processing was performed using Zeiss application software. Leaf samples from Arabidopsis plants were collected at 6 hpi (hours post infiltration) to examine the induction of selected differentially expressed genes in response to the *Pst*DC3000 challenge using the qRT-PCR assay. The leaves infiltrated with 10 mM MgCl_2_ were taken as control. Actin and ubiquitin were used as endogenous controls for gene normalization.

### Statistical analysis and R packages used

The experiments in the study were performed in two independent biological replicates, each with three technical repeats. The data shown in the study are mean ± SD. Student t-test was performed using instat.exe version 3.0 software to measure the degree of significance with a P-value ≤ 0.05 and denoted by an asterisk above the bar graph in the figures. P-values ≤ 0.001 are designated with three asterisks while p values ≤ 0.0001are designated with four asterisks. For GSEA Go enrichment analysis, R cluster Profiler and enrich plot package were used. For the ridge plot, the r ggridges package was used. The Tidyverse package was used to make a volcano, dot plot, and bar plots. For heatmap, the pheatmap package was used.

## Results

### *AtERF60* is induced under salt, dehydration, ABA and SA treatments

The *AtERF60* CDS is 819 bp long and encodes a 273 amino acid polypeptide. The alignment of complete ORF with its amino acid sequence has shown that it has 58 amino acids conserved AP2/ERF DNA binding domain (DBD) (Fig. S1). The phylogenetic homology of *AtERF60* was investigated using MEGAXI software with the neighbour-joining method and 1000 bootstrap repetitions, and it was discovered that it is closely related to Arabidopsis *ERF056*. (Fig. 1a). Chromosomal location and mutant information used in the study are depicted in (Fig.1 b). To study *AtERF60* under various conditions, we checked the induction of *AtERF60* in response to abscisic acid (ABA), salt, dehydration, (SA) salicylic acid, wounding and methyl jasmonate (MeJA) treatments in *Arabidopsis thaliana* Col-0 plants. We observed that *AtERF60* is induced early after 1 hour of ABA treatment. *AtERF60* showed maximum induction after 3 hours of salt and dehydration treatment. Transcripts of *AtERF60* showed a similar level of expression after 1, 3 and 5 hours of SA treatment (Fig. 1c). Additionally, *AtERF60* is not induced in response to MeJA and wounding treatment as compared to the untreated control WT Col-0 (Fig. 1c).

**Figure 1.**
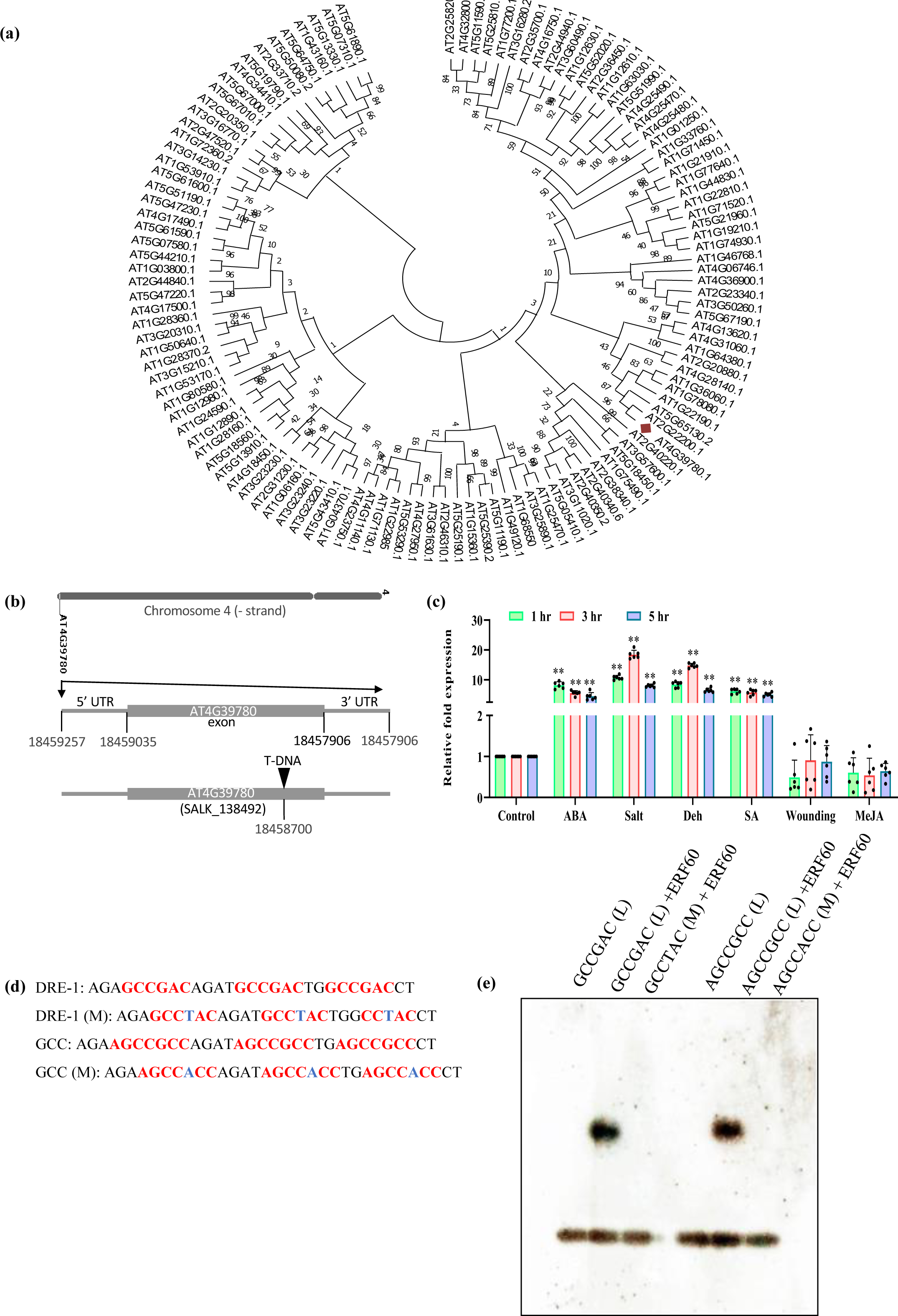
Phylogeny, expression analysis, and Electrophoretic mobility shift assay (EMSA) of *AtERF60*. **(a)** The protein sequences of 122 AP2/ERF family members from Arabidopsis thaliana were retrieved from the TAIR database. The amino acid sequences were aligned by clustalW and the phylogenetic tree was constructed using MEGAXI by neighbour-joining (NJ) method and 1000 bootstrap replicates. **(b)** Gene structure of *AtERF60*, its chromosome location and T-DNA insertion line**. (c)** Relative expression of *AtERF60* transcript in response to ABA, salt, dehydration, SA, wounding, and MeJA treatment after 1, 3, and 5 h. Relative expression of transcripts was calculated by taking untreated plant samples as a control (WT). Actin and ubiquitin were used as endogenous controls for gene normalisation. Error bars indicate mean ± SD. Student’s t-test, **, P < 0.01. **(d)** Probes containing DRE (GCCGAC) and GCC-box (AGCCGCC) *cis*-elements were designed to study the DNA-protein interaction. The desired *cis*-elements are red, while the binding site carrying mutations are marked blue. **(e)** EMSA of *AtERF60* showed that it interacts specifically with DRE-1 and GCC regulatory *cis*-elements (M-mutated/substituted and L-DIG-labelled).

### Recombinant AtERF60 interacts with DRE/GCC box *cis*-elements

The recombinant GST-tagged-AtERF60 was induced and purified in the bacterial expression system (Fig. S2b, c). An expected molecular mass of 56.0 kDa along with GST-tagged-AtERF60 was observed and separated using SDS-PAGE (Fig. S2b). To study the DNA binding property of *AtERF60*, we performed EMSA of recombinant AtERF60 with DRE-1 and GCC-box *cis*-elements. Specific probes were designed for DRE-1 (GCCGAC) and GCC-box (AGCCGCC) *cis*-element along with their mutated forms containing a single nucleotide substitution (Fig. 1d). We found that the recombinant AtERF60 interacts with DRE-1 and GCC-box *cis*-elements, while it does not bind with its mutated probe carrying a change of single nucleotide (Fig. 1e). The EMSA demonstrate that *AtERF60* TF interacts specifically with DRE-1 and GCC-box *cis*-elements and might be involved in the regulation of downstream target genes.

### *AtERF60* regulates basal resistance response in Arabidopsis

ABR1 is a functionally important molecule that interacts with different *Pseudomonas syringae* effectors (Schreiber *et al*., 2021). We sought to test our idea that *AtERF60* regulates the central key component *ABR1* and modulates plant response to biotic stress. We generated independent homozygous lines by constitutively overexpressing *AtERF60* under *CaMV 35S* to investigate its role in response to biotic stress (Fig S3 a, b, c d) and validated the overexpression lines (Fig 2a). Additionally, separate homozygous lines were generated by complementing the *erf60* mutant lines (Fig 2a). The *AtERF60-*OX, *erf60* mutant, and *AtERF60* complemented lines were infected with *Pseudomonas syringae* pv. *tomato* (*Pst*) strain DC3000 to examine the bacterial pathogen response. The *AtERF60*-OX lines exhibited reduced chlorosis and necrosis at 4 days of post-inoculation (dpi), whereas *erf60* mutant lines displayed increased disease symptoms as compared to the Col-0 plants. Furthermore, the *erf60* mutant lines supported a significantly higher population of bacterial *Pst*DC3000 than the Col-0 plants. The bacterial population in the *AtERF60-* OX lines was reduced by 0.93 log (CFU/cm^2^) at 2 dpi while it was reduced by 1.34 log (CFU/cm^2^), after 4 dpi (Fig. 2c). The *erf60* lines, on the other hand, have a 0.61 log (CFU/cm2) and 0.70 log (CFU/cm2) increase in bacterial population at 2 and 4 dpi, respectively (Fig. 2c). A significant reduction in electrolyte leakage was observed in *AtERF60-*OX lines (∼16.04 %) compared to the Col-0 plants at 4 dpi with *Pst*DC3000 (Fig. 2d). Contrary to that, electrolyte leakage was found to be significantly higher in *erf60* lines (∼ 29.43 %) compared to the Col-0 (Fig. 2d) plants. No significant difference in the bacterial population and electrolyte leakage was observed in the *AtERF60* complemented lines compared to Col-0. Further, we used confocal microscopy to observe the colonization of *Pst*DC3000 in Arabidopsis plants using the GFP-tagged *Pst*DC3000 strain. In vivo imaging revealed the formation of the distinct microcolonies of GFP-tagged *Pst*DC3000 that were evenly dispersed in the intercellular spaces of leaves of Col-0 plants. We observed reduced colonization of GFP-tagged *Pst*DC3000 in *AtERF60-* OX lines while *erf60* lines displayed enhanced colonization of bacterial pathogen (Fig. 2b). Together, these results suggest that mutation in the *AtERF60* gene causes an increase in susceptibility of Arabidopsis towards bacterial pathogen, *Pst*DC3000.

**Figure 2.**
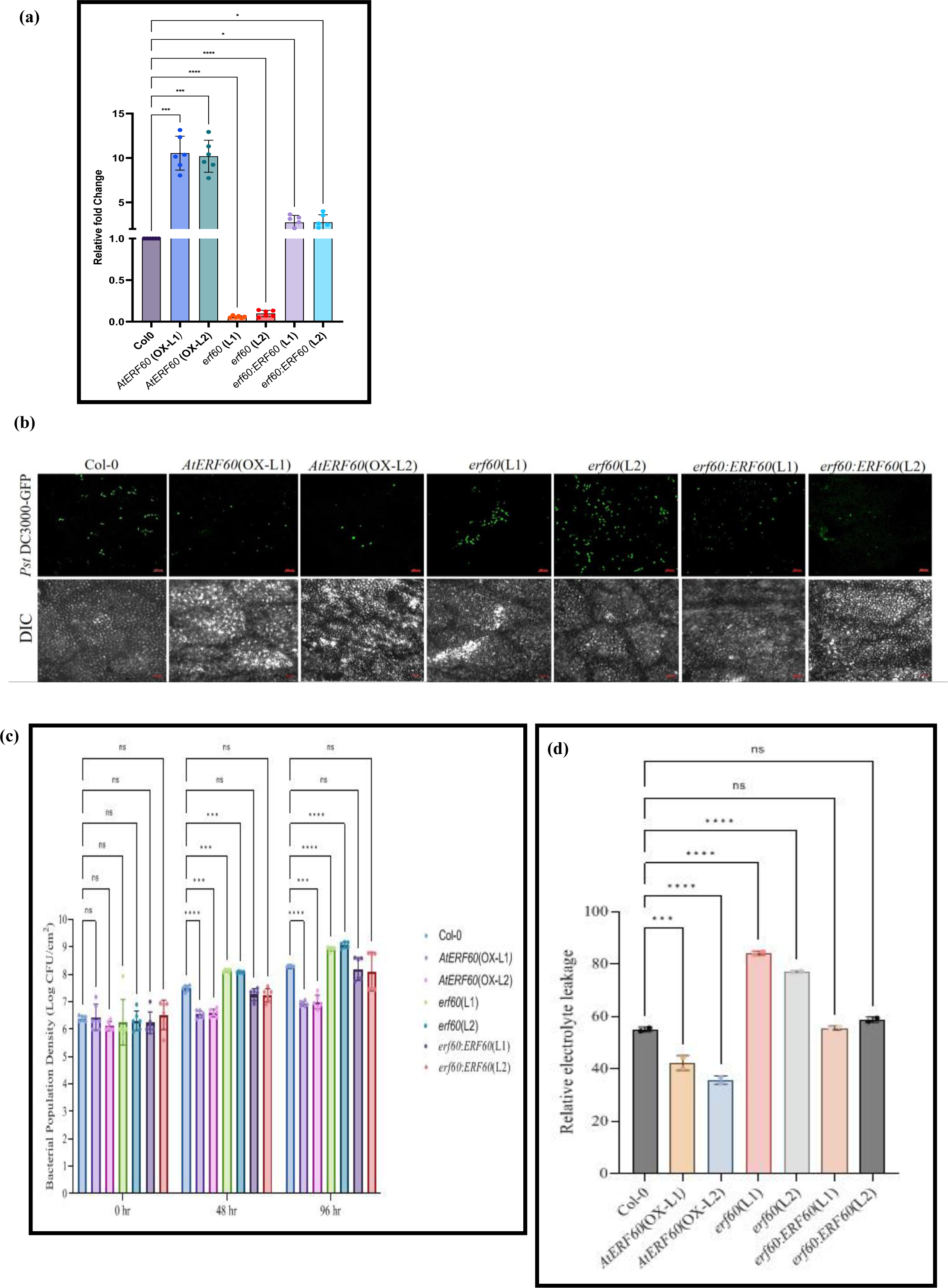
Evaluation of *Pst*DC3000 colonization in *AtERF60* overexpressing, mutant and complemented lines. **(a)** Relative expression of *AtERF60* transcripts in overexpression lines (OX), *erf60* mutant lines and *AtERF60* complementation lines compared to Col-0 plants. **(b)** Differential interference contrast (DIC) and confocal images of leaves of WT, *AtERF60-OX* and *erf60* mutant lines with GFP-tagged *Pst*DC3000 at 2 dpi. GFP fluorescence (green) was acquired using 488 nm excitation and 493-598 nm emission. Images of DIC and confocal microscopy (scale bar, 100 µm). **(c)** Bacterial growth estimation in leaves of WT, *AtERF60-*OX, *erf60* mutant and *AtERF60* complemented lines inoculated with *Pst*DC3000 at 0, 2, and 4 dpi. 2-way anova was used for the analysis. **(d)** Electrolyte leakage from leaves of WT, *AtERF60-*OE *erf60* mutant and *AtERF60* complemented lines inoculated with *Pst*DC3000 at 4 dpi. Electrolyte leakage values are given as the percentage of total ions.1-way anova was used for the analysis. Actin and ubiquitin were used as endogenous controls for gene normalization. The error-bars represents SE, P-values ≤ 0.001 are designated with three asterisks while P values ≤ 0.0001are designated with four asterisks.

### Loss of *AtERF60* function affects multiple gene expression

To understand the regulatory role of *AtERF60* in Arabidopsis, we performed a microarray analysis of *AtERF60*-OX and *erf60* mutant lines under control conditions in two independent biological replicates (Table S1). We observed a significant differential expression gene (DEG) change in mutants and OX lines. There are 1527 DEGs upregulated and 1050 DEGs downregulated in OX lines with p-value >0.05 and 2-fold change increase/decrease, while 1226 DEGs are upregulated and 773 are down-regulated in mutant lines. (Fig. 3a, b). Further, these DEGs are categorised into function GO categories. The categories are differentiated based on whether they are activated or suppressed in biological processes, cellular components and molecular function. The top 10 enriched categories in OX and mutant lines are shown (Fig. 3c, e). Photosynthesis-associated pathways are seen enriched in, and protein quality control pathways are suppressed in OX lines while Sugar pathways are enriched in mutant and transport pathways are suppressed. To interpret up/down-regulated pathways grouped by gene set, density plots are generated using the frequency of fold change values per gene within each set of top enriched categories in OX and mutant lines (Fig. 3d, f). We further validated the expression of 13-above identified genes using qRT-PCR analysis in both *AtERF60*-OX *erf60* and its complemented lines as compared to the Col-0 (Fig. 4). We found that most of the genes encoding *COMPROMIZED RECOGNITION OF TCV1 (CRT1), ABR1, Peroxidase superfamily protein, Aspartyl protease family protein, O-glucosyl hydrolases family protein, Alcohol dehydrogenase 1 (ADH1), Acyl-CoA synthetase 5 (ACS5),* and *EIDI like 3*, were significantly upregulated in the *erf60* mutant background as compared to the Col-0. Complementation of *AtERF60* in a mutant background restored the increased expression of these downregulated transcripts to their wild type levels, indicating that *AtERF60* suppresses the expression of these genes (Fig 4). The microarray and qRT-PCR analysis revealed that *AtERF60* might be involved in downregulating these target genes directly or indirectly by binding with their promoter *cis*-elements. Therefore, we screened out the individual promoters and searched for probable *cis*-elements (Supplemental File 1).

**Figure 3.**
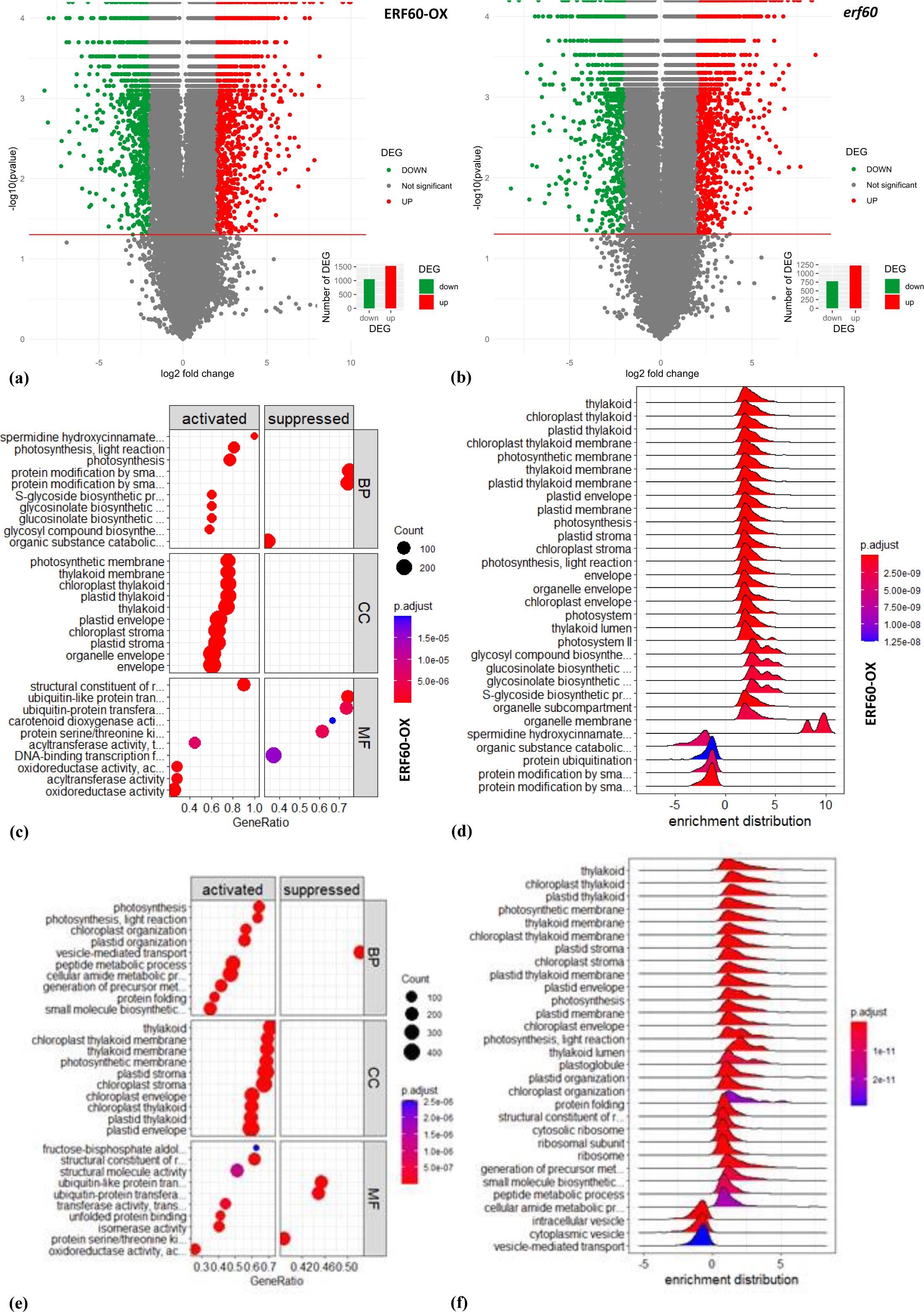
Microarray analysis of *AtERF60-*OX and *erf60* mutant plants under controlled conditions. **(a)** For the *AtERF60* overexpression lines (OX), univariate analysis of DEGs in volcano plot with negative logarithm of the p value on the y-axis and logarithm of the fold change on the x-axis. Red dots show upregulated genes with fold change ≥ 2, and blue dots show down-regulated genes with fold change ≤ 2. The red threshold line shows a p-value of 0.05. The grey dots are non-significant DEGs. The subplot shows the number of up-and-down-regulated genes. **(b)** For the mutant line, volcano plot and bar plot of up-and down-regulated DEGs. **(c)** Enrichment plot (dot plot) of OX lines with GO categories up and down-regulated in biological processes (BP), cellular component (CC) and molecular function (MF). **(d)** Ridge plot (density plot) of OX lines summarizing the “core enriched” genes for each gene set via (log2) fold-change. The colour describes the adjusted p-value listed in the colour bar, notifying whether an enriched term is significant. **(e)** Enrichment dot plot and **(f)** ridge plot of mutant lines showing GO categories and core enriched genes.

**Figure 4.**
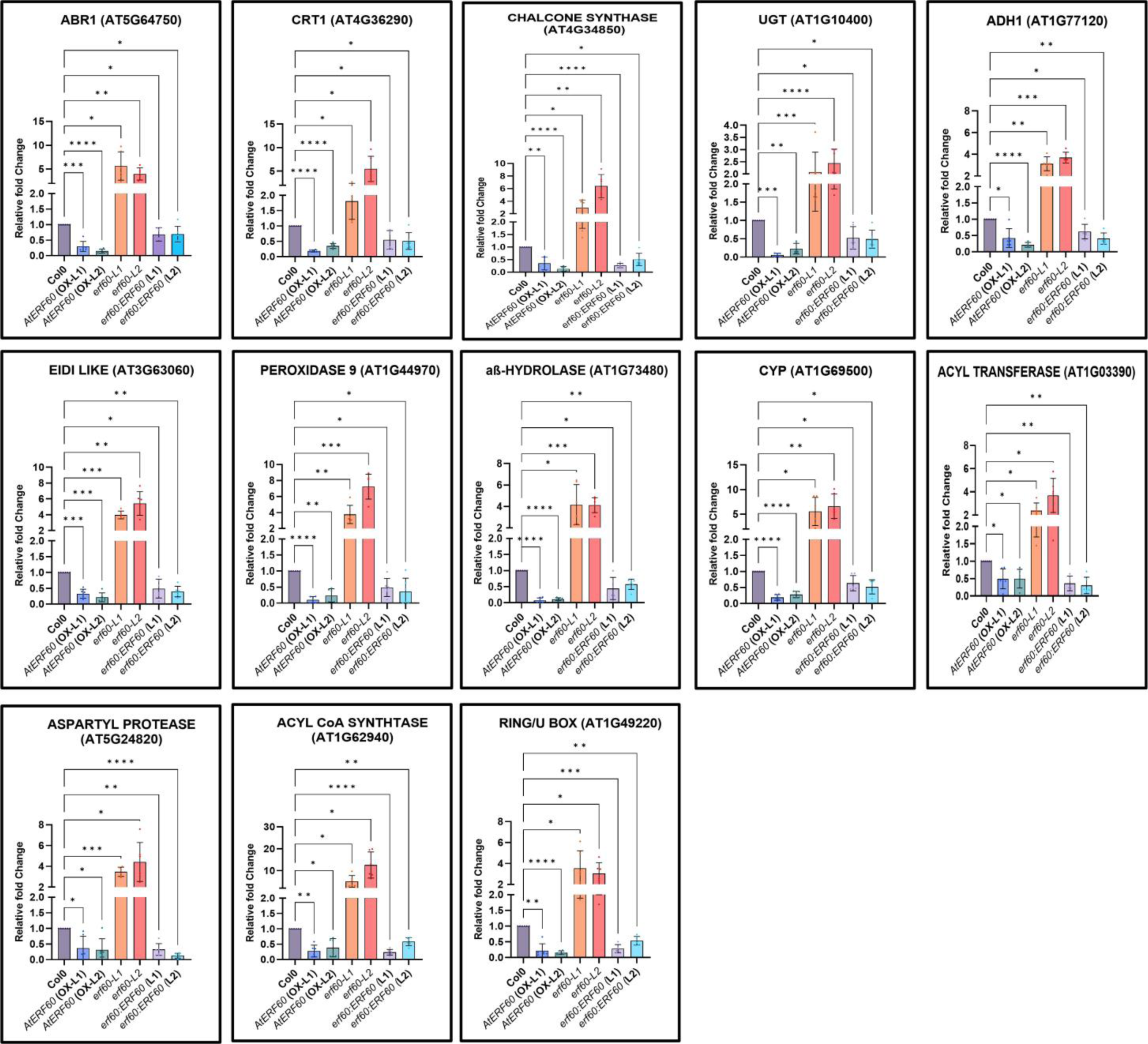
Validation of differentially expressed target genes between *AtERF60-*OX, *erf60* mutant and complementation lines through qRT-PCR. Relative expression of differential target genes as determined by qRT-PCR in the *AtERF60-*OX, *erf60* mutant and *AtERF60* complemented lines compared to the WT. 13 differentially expressed genes were selected for validation through qRT-PCR. Error bars indicate mean ± SD. P-values ≤ 0.001 are designated with three asterisks while P values ≤ 0.0001are designated with four asterisks. Actin and ubiquitin were used as endogenous controls for gene normalization.

### *AtERF60* interacts with *ABR1* promoter

Sequence analysis revealed the presence of dehydration (A/GCCGAC) and ABA-responsive *cis*-elements (ACGTC/G) in the *ABR1* promoter. Based on the presence of these core *cis*-elements in the *ABR1* promoter, we studied the interaction of *AtERF60* with the *ABR1* promoter using EMSA. Probes specific to DRE and ABRE *cis*-elements were designed constituting the promoter region of ABR1 along with their mutated counterpart with a change in a single nucleotide (Fig. 5a). It was observed that *AtERF60* interacts with both DRE-1 and DRE-2 *cis*-elements (Fig. 5b). At the same time, it is not able to produce gel shift with the ABRE *cis*-elements and its counterpart with a single nucleotide mutation (Data not shown). This showed that the binding of *AtERF60* is specific to A/GCCGAC *cis*-elements and is completely abolished when the sequence-specific nucleotide is mutated.

**Figure 5.**
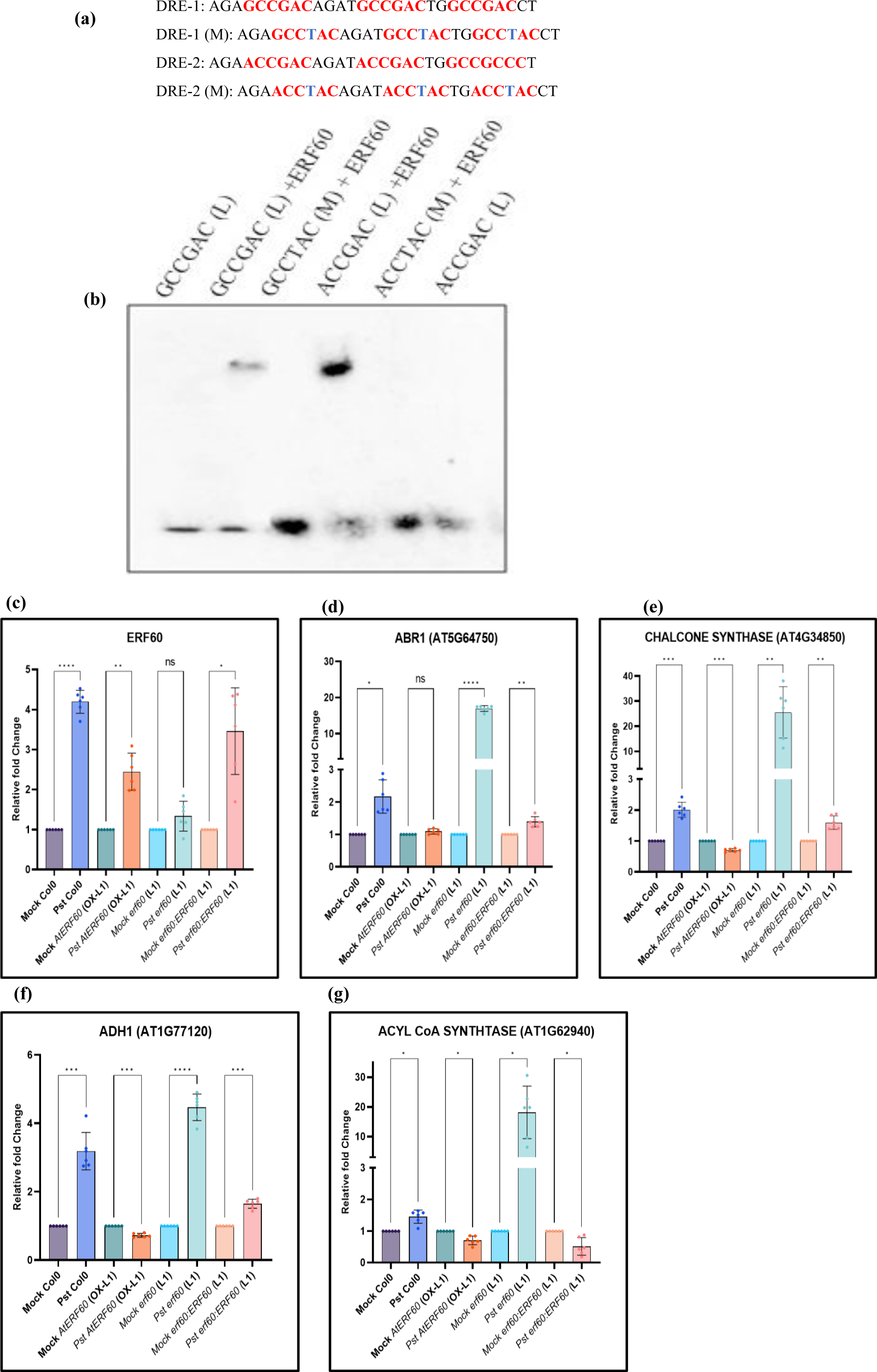
EMSA analysis to determine binding ability of AtERF60 with ABR1 promoter region. **(a)** Probes containing DRE1/2 (A/GCCGAC) *cis*-elements were designed to study the DNA-protein interaction. The desired *cis*-elements are red, while the binding site containing mutations is blue. **(b)** EMSA of *AtERF60* showed that it interacts with the regulatory DRE1 and DRE2 cis-elements in the *ABR1* promoter. In contrast, it does not interact with the ABRE1/2 *cis*-elements (M-mutated/substituted and L-DIG-labelled). **Validation of differentially expressed target genes between AtERF60-OX, erf60 mutant and complementation lines after 6-hour post inocoulation of PstDC3000**. qRT-PCR analysis of **(c)** *ERF60*, **(d)** *ABR1*, **(e)** *chalcone synthase*, **(f)** *ADH1* and **(g)** *ACS* genes in Col-0, OX and mutant lines following infection with *Pst* DC3000 after 6 hours post-inoculation (hpi) relative to Col-0 infiltrated with 10mM MgCl_2_ (mock). Error bars indicate mean ± SD. P-values ≤ 0.001 are designated with three asterisks while P values ≤ 0.0001are designated with four asterisks. Actin and ubiquitin were used as endogenous controls for gene normalization.

We examined the expression of *AtERF60* and *ABR1* genes in Arabidopsis plants following infection with *Pst*DC3000. The expression of the *AtERF60* and *ABR1* genes are induced in Col-0 in response to *Pst*DC3000 infection as observed at 6 hours post-inoculation (hpi) (Fig 5c, d). On the other hand, *ADH1, ACS, CHS*, and *ABR1* genes show induced expression in the *erf60* mutant plants, followed by a *PstD*C3000 infection of 6 hours. The expressions of these transcripts were downregulated in the *AtERF60*-OX lines after 6 hours of *Pst*DC300 infection (Fig 5d-g). When *AtERF60* is complemented into *erf60* mutant lines, it causes a reversal in the expression of these transcripts to the Col-0 level. Moreover, after being challenged with 6 hours of *Pst*DC3000, the *erf60* mutant plants show a considerable increase in *ABR1* expression, while the *AtERF60*-OX plants show a significant downregulation. (Fig. 5c-g). Thus, our data suggest a negative regulation of *ABR1* by *AtERF60* under biotic stress response.

## Discussion

AP2/ERFs regulate various downstream signalling cascades by interacting with different *cis*-elements of target gene promoters and regulating multiple responses (Phukan *et al*., 2018). In this study, we have identified a transcription factor, *AtERF60*, induced in response to drought conditions, salt stress, ABA/SA treatment, and bacterial pathogen *Pst*DC3000 infection. Induction of the *AtERF60* in response to ABA and SA suggests that the *AtERF60* might regulate downstream responses in an ABA/SA-dependent manner. In Arabidopsis, *ERF53* (Hsieh *et al*., 2013), *RAP2.6* (Zhu *et al*., 2010), and *ERF-VIIs* (Yao *et al*., 2017) are induced in response to ABA to upregulate the genes containing DRE/ABRE *cis*-elements. There are limited numbers of AP2/ERFs reported to be involved in response to SA treatment (Xie *et al*., 2019). The potential of AP2/ERFs to counter different signals and regulate multiple stresses facilitates them to make a highly complex stress regulatory network. Some AP2/ERFs are induced at earlier stages, while others are induced upon prolonged stress, which suggests that their function might be influenced mutually (Van den Broeck *et al*., 2017).

The diversity of AP2/ERFs in response to various stresses depends on the flexibility of the AP2 domain that implements the binding to different *cis*-elements such as DRE/CRT and GCC-box elements (Huang *et al*., 2008, Cheng *et al*. 2013, Franco-Zorrilla *et al*., 2014, Catinot *et al*., 2015). Our previous study conclusively demonstrated that AP2/ERF TFs such as *PsAP2* and *MaRAP2-4* from the *Opium poppy* and *Mentha arvensis*, respectively, bind with both GCC and DRE box *cis*-elements to modulate biotic and abiotic stress response in transgenic plants (Mishra *et al*., 2015; Phukan *et al*., 2018). We observed that it interacts with the core *cis*-element of DRE (GCCGAC), and the binding was abolished when we used the mutated version by changing the 4^th^ nucleotide G to T (GCC**T**AC). We also observed that *AtERF60* interacts with the core GCC box element GCCGCC, and mutation of the probe into GCCACC abolished its interaction. These results suggest that CCGCC and CCGAC are the core sequences preferred for *AtERF60* to interact. If we mutate G of CCGCC to CCACC, the binding was abolished. We find that the mutation in the last two CC to AC for the DRE sequence does not affect its binding ability. Some ERFs, including *ERF71*, *ERF4*, and *ERF1*, are also reported to interact with both DRE and GCC elements in Arabidopsis (Lee *et al*., 2015; Xie *et al*., 2019). The ability of *AtERF60* to interact specifically with DRE and GCC further supports its probable involvement in stress response.

Additional insight into *AtERF60* function in biotic stress response was gained by processing and analysing the microarray and expression data in both the overexpression and mutant background. Microarray analysis, promoter binding activity, and phenotypic studies suggest the regulatory role of *AtERF60* in Arabidopsis. The fold expressions of the majority of differentially expressed transcripts are significantly higher in the *erf60* mutant background under control. Additionally, we identified the upregulation of target genes in the *erf60* mutant plants following *Pst*DC3000 inoculation, including *ABR1, chalcone synthase gene*, *Acyl CoA synthase 5,* and *ADH1*.

In this study we discovered that, the mutation in *AtERF60* enhanced susceptibility against bacterial pathogen *Pst*DC3000. In contrast, *AtERF60*-OX plants exhibited reduced susceptibility to *Pst*DC3000. This work identifies *AtERF60* as the pathogen-responsive gene which downregulates *ABR1,* which contributes to plant immunity by interacting with *Pseudomonas syringae* effector molecule HopZ1a and other different effectors (Schreiber *et al*., 2021). Increased expression of *ABR1* in the *erf60* mutant plants in response to the pathogen infection suggests that *AtERF60* downregultes the *ABR1* gene expression during biotic stress. Under similar conditions, *AtERF60*-OX plants show a suppression of *ABR1* expression. Our results suggest that *AtERF60* plays an important role in plant bacterial-pathogen interaction via downregulating *ABR1* expression under the control conditions. The *AtERF60*-OX lines exhibited reduced susceptibility, and its functional mutant showed an enhanced susceptibility response. Our results conclude that mutation in *AtERF60* makes susceptibility hub *ABR1* constitutively active, leading to susceptible phenotype towards bacterial pathogen *Pst*DC3000.

## Acknowledgements

The authors acknowledge the Director, CSIR-CIMAP, for providing the necessary facilities. Authors acknowledge ABRC for providing the mutant seeds of *AtERF60*. We also acknowledge Genotypic for performing the microarray analysis of overexpression and mutant background of *AtERF60*.

## Author contributions

AJ has generated the complementation lines, contributed to the bacterial pathogen infiltration experiments and CFU analysis, and performed electrolyte leakage; UJP has cloned the AtERF60 gene made the over-expression lines and screened the *erf60* mutant lines, giving samples for microarray analysis and contributed to analysis of figure 3. NS has contributed to the bacterial pathogen infiltration experiments and CFU analysis and electrolyte leakage, maintained the seed stock and plating of seeds. GSJ has performed EMSA experiments. SK has completed validation of q-RT-PCR of *AtERF60-OX*, *erf60* and its complementation lines under the control condition and after bacterial infiltration. DP has contributed to bacterial CFU analysis. VT has contributed to planning experiments related to mutant background and procuring of the mutant background of *AtERF60*. AP has supervised the bacterial colony infiltration. AJ, SK, NS, and RKS has analyzed the result, RKS has supervised the work, planned the experiments and edited the manuscript. AJ, SK, NS, and RKS have finalized the draft manuscript.

## Data Availability Statement

*ERF60*-AT4G39780, *ADH 1*-AT1G77120, *ACOS5*-AT1G62940, *Chalcone and stilbene synthase*-AT4G34850, *CYP450 family protein*-AT1G69500, *Aspartyl protease family protein*-AT5G24820, *Actin 1*-AT2G37620, *Acyl-transferase family protein*-AT1G03390, *EID1-like 3*-AT3G63060, *ABR1*-AT5G64750, *CRT1*-AT4G36290, *alpha/beta-Hydrolases superfamily protein*-AT1G73480, *O-Glycosyl hydrolases family 17 protein*-AT4G14080, *Myb-like HTH transcriptional regulator family protein*-AT1G18960, *UDP-Glycosyltransferase superfamily protein*-AT1G10400, *RING/U-box superfamily protein-* AT1G49220, Microarray data deposited with the accession number GSE179600.

## Supplementary data

**Figure S1. Nucleotide and amino acid alignment of *AtERF60***. Complete cDNA (1224 bp) carrying 819 bp of ORF. Red represents the amino acid sequence, and blue represents the AP2/ERF DNA binding domain. *AtERF60* has a 145 bp 5’ UTR and 259 bp of 3’ UTR. It encodes a protein of 273 amino acids having a molecular weight of 30 kDa.

**Figure S2. Protein induction and purification of *AtERF60*. (a)** Confirmation of positive clones of *AtERF60* by PCR and restriction digestion in bacterial expression vector pGEX-4-T2. **(b)** The recombinant AtERF60 protein was induced with different concentrations (mM) of IPTG at 37^0^C. **(c)** The AtERF60-GST fusion protein was purified from the *E. coli* BL21 (DE3) strain following induction for 5 hours at 37^0^C with 0.3 mM IPTG. An affinity-purified recombinant protein with a molecular weight of 56.0 kDa was segregated on SDS-PAGE.

**Figure S3. Cloning and validation of transgenic lines of *AtERF60*. (a)** Confirmation of positive clones of *AtERF60* by PCR and restriction digestion in plant expression vector pBI121. **(b)** Selection of transgenic lines on kanamycin-supplemented half MS media. **(c)** PCR confirmation of transgenic lines. Actin was used as an internal control. **(d)** Genomic DNA PCR of transgenic lines with NPT II (KanR) primers.

**Figure S4. Mutant screening of *AtERF60* lines.** Genomic DNA PCR of mutant *AtERF60* Salk lines. LB-left border, RP-right primer, LP-left primer. The resulting amplification using primers specific LB and RP showed mutant homozygous lines.

**Table S1.** Microarray data showing the list of up and downregulated genes obtained.

**Table S2.** List of genes identified from microarray analysis with significant p-value (<0.05).

**Table S3.** List of primers used in the study.

**Supplementary file 1**. List of selected Arabidopsis promoter sequences. The cis-elements are marked with green colors.

