## Supplemental Figures for "*AtERF60,* negatively regulates *ABR1* to modulate basal resistance response in Arabidopsis"

**Fig. S1**

```
aaaaaacaaatatttagaaacaaaaaatgctataattcttccttttcttgttctttcag
agatttgatttctattcaatcgactaagagggtgtgttttaatccctctctgttttaa
ttttgtttcctagtctttaaagatcc
atggcagccatagatatgttcaatagcaacacagatccttttcaagaagagctcatgaaa
 M  A  A  I  D  M  F  N  S  N  T  D  P  F  Q  E  E  L  M  K
gcacttcaaccttataccaccaacactgattcttcttctcctacgtattcaaacacagtc
 A  L  Q  P  Y  T  T  N  T  D  S  S  S  P  T  Y  S  N  T  V
ttcggtttcaatcaaaccacatctctcggtctaaaccagctcacaccttaccaaatccac
 F  G  F  N  Q  T  T  S  L  G  L  N  Q  L  T  P  Y  Q  I  H
caaatccaaaaccagcttaaccagagacgtaacataatctctccaaatctagccccaaag
 Q  I  Q  N  Q  L  N  Q  R  R  N  I  I  S  P  N  L  A  P  K
cctgtcccaatgaagaacatgaccgctcagaaactctatagaggagttagacaaaggcac
 P  V  P  M  K  N  M  T  A  Q  K  L  Y  R  G  V  R  Q  R  H
tggggaaaatgggtagctgagatccgtttacccaagaaccggacccgactctggcttgga
 W  G  K  W  V  A  E  I  R  L  P  K  N  R  T  R  L  W  L  G
actttcgacacagctgaagaagcagccatggcttatgacctagctgcttacaagctaaga
 T  F  D  T  A  E  E  A  A  M  A  Y  D  L  A  A  Y  K  L  R
ggcgagttcgcgagacttaatttcccacagttcagacacgaggatggatactacggagga
 G  E  F  A  R  L  N  F  P  Q  F  R  H  E  D  G  Y  Y  G  G
ggtagctgtttcaatcctcttcattcctctgtcgacgcaaagctccaagagatttgtcag
 G  S  C  F  N  P  L  H  S  S  V  D  A  K  L  Q  E  I  C  Q
agcttgagaaaaacagaggatattgacctcccctgttctgaaacagagcttttcccgcca
 S  L  R  K  T  E  D  I  D  L  P  C  S  E  T  E  L  F  P  P
aaaacagagtatcaagaaagtgaatatgggttcttgagatctgatgagaattcgttttca
 K  T  E  Y  Q  E  S  E  Y  G  F  L  R  S  D  E  N  S  F  S
gatgagtctcatgtggaatcttcttcgccggaatctggtattactacgttcttggacttt
 D  E  S  H  V  E  S  S  S  P  E  S  G  I  T  T  F  L  D  F
tcggattctggatttgatgagattgggagtttcgggctggagaagtttccttctgtggag
 S  D  S  G  F  D  E  I  G  S  F  G  L  E  K  F  P  S  V  E
attgattgggatgcgattagcaaattgtccgaatcttaa
 I  D  W  D  A  I  S  K  L  S  E  S  -
acaaagcaaagagaagacttttcttttaggagtttgtctttcaatttcagtgtcttata
ttaatctctctgcaactgaaattttaacagttgcggagagaatcgtctctagggtttgt
ttctcttcctccatgttttggtctgactggtttatgtctttttttttttttgaactttc
aaaaacctttgtcattgaccaatcggagaagtttccatgtattctctcatctttaaattt
ctaatgaagaatgtaaatt
```

**Fig. S2**

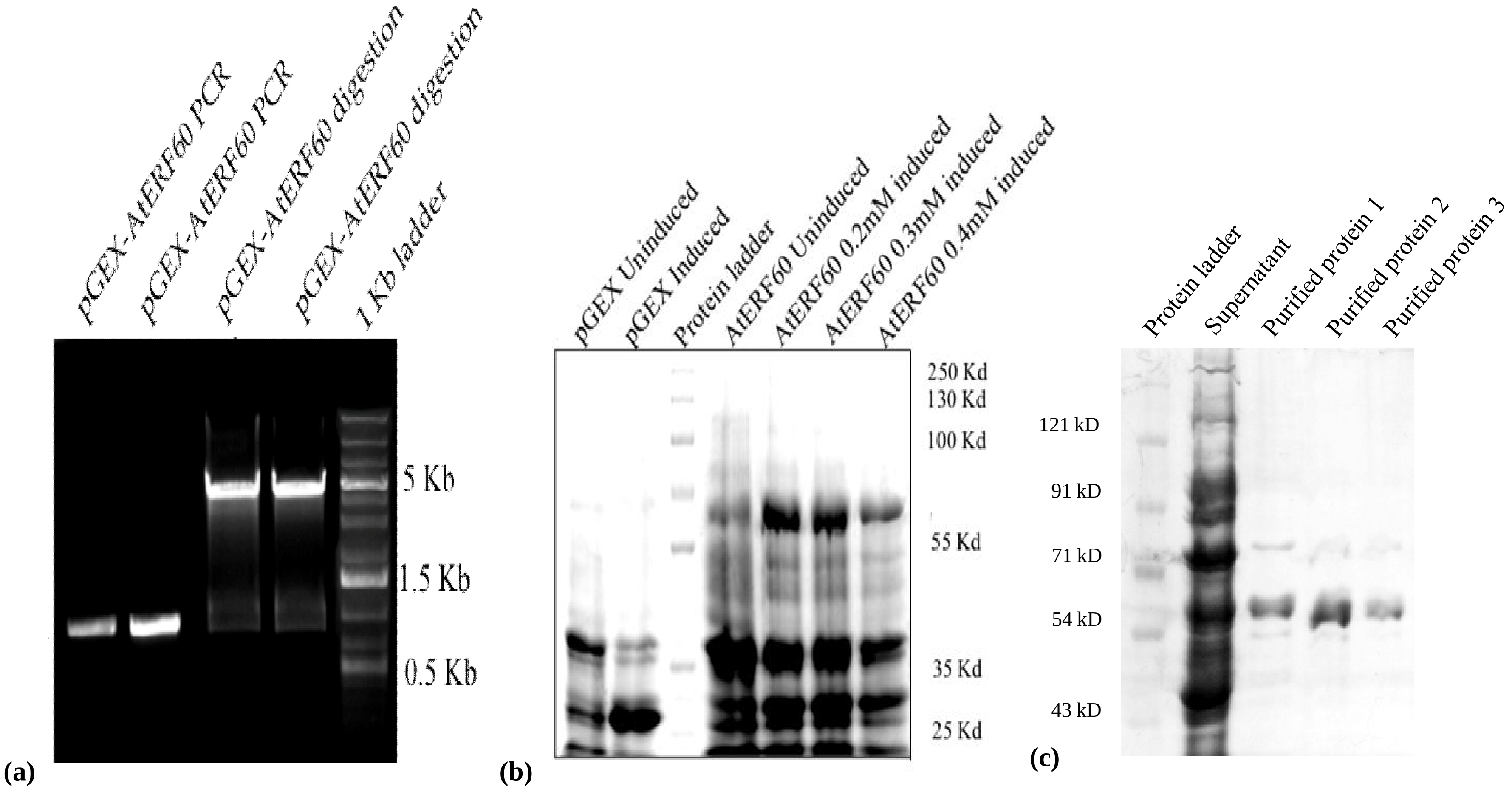
pGEX-AtERF60 PCR
pGEX-AtERF60 PCR
pGEX-AtERF60 digestion
pGEX-AtERF60 digestion
1 Kb ladder
5 Kb
1.5 Kb
0.5 Kb
(a)
pGEX Uninduced
pGEX Induced
Protein ladder
AtERF60 Uninduced
AtERF60 0.2mM induced
AtERF60 0.3mM induced
AtERF60 0.4mM induced
250 Kd
130 Kd
100 Kd
55 Kd
35 Kd
25 Kd
(b)
Protein ladder
Supernatant
Purified protein 1
Purified protein 2
Purified protein 3
121 kD
91 kD
71 kD
54 kD
43 kD
(c)

**Figure S3**

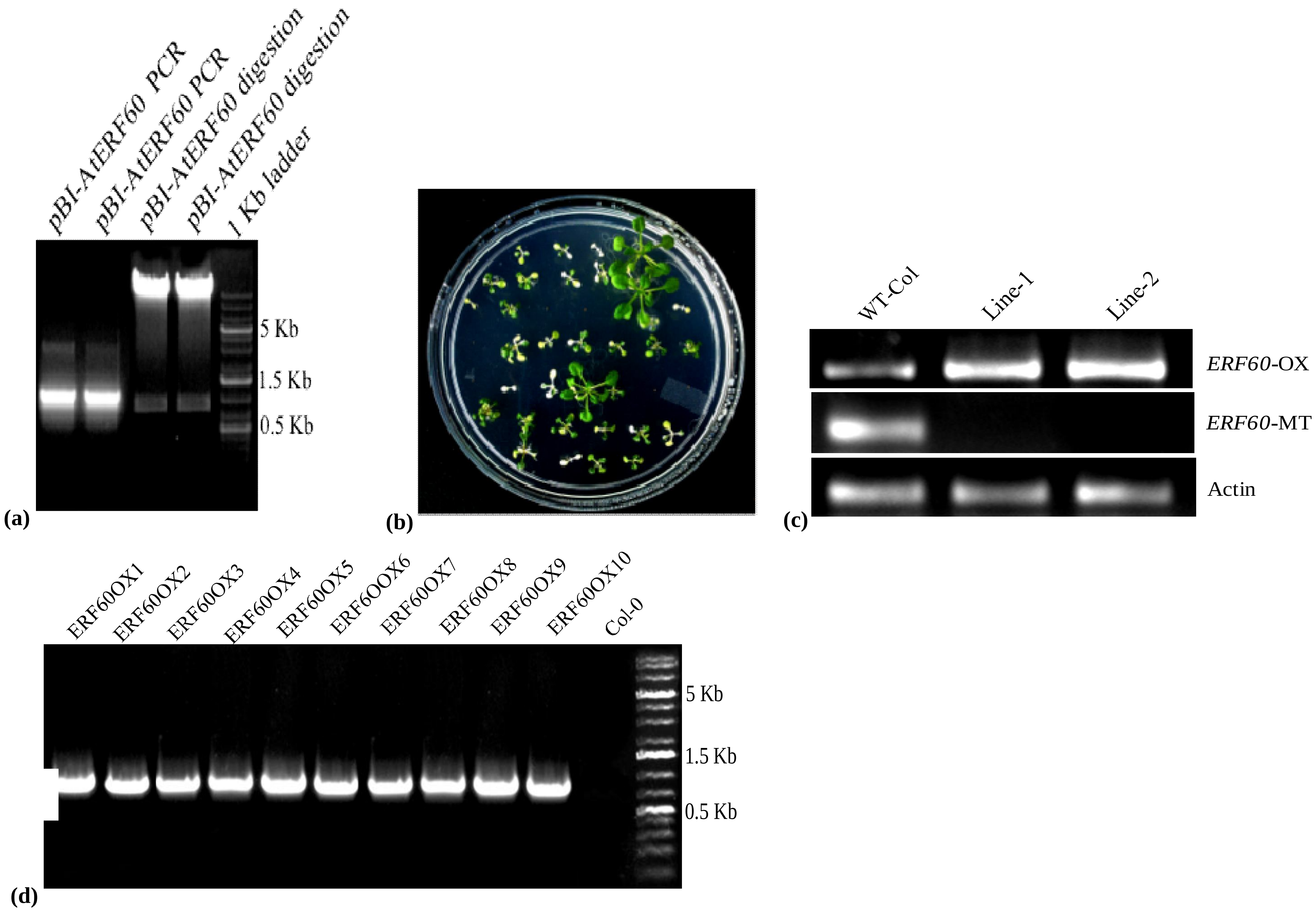
pBI-AtERF60 PCR
pBI-AtERF60 PCR
pBI-AtERF60 digestion
pBI-AtERF60 digestion
1 Kb ladder
5 Kb
1.5 Kb
0.5 Kb
(a)
(b)
WT-Col
Line-1
Line-2
ERF60-OX
ERF60-MT
Actin
(c)
ERF60OX1
ERF60OX2
ERF60OX3
ERF60OX4
ERF60OX5
ERF60OX6
ERF60OX7
ERF60OX8
ERF60OX9
ERF60OX10
Col-0
5 Kb
1.5 Kb
0.5 Kb
(d)

**Figure S4**

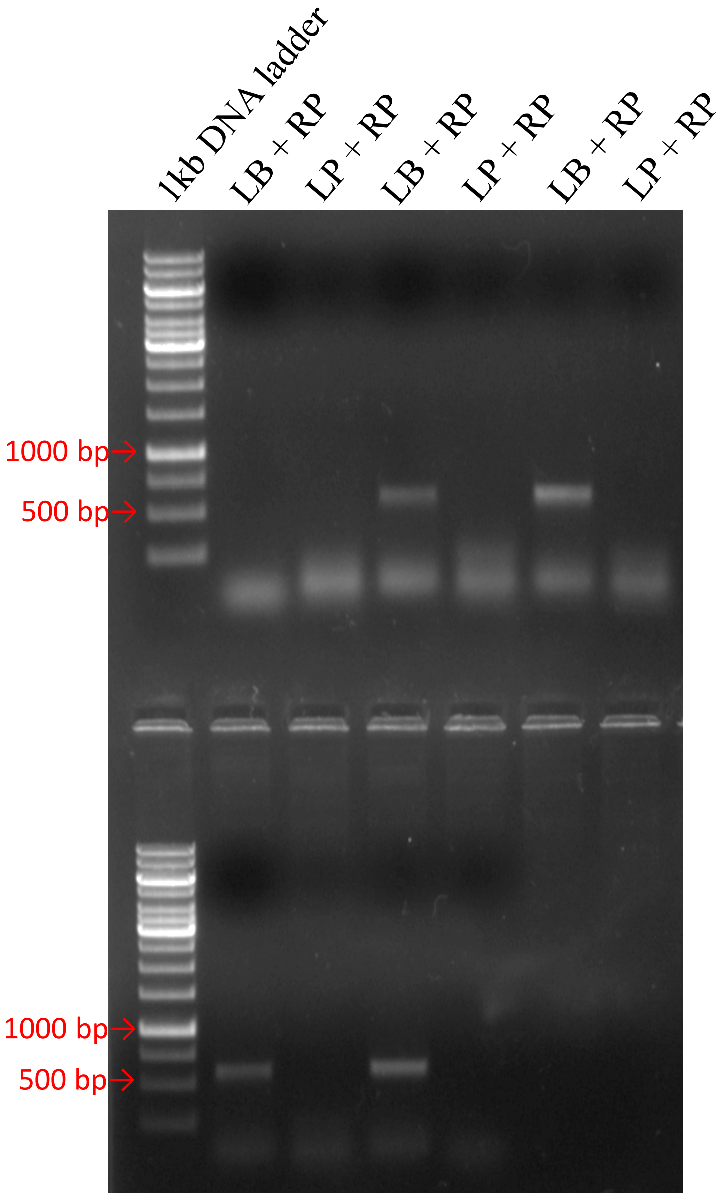
1kb DNA ladder
LB + RP
LP + RP
LB + RP
LP + RP
LB + RP
LP + RP
1000 bp
500 bp
1000 bp
500 bp
